# De novo annotation of centromere with centroAnno

**DOI:** 10.1101/2025.02.19.639205

**Authors:** Junhai Qi, Junchi Ma, Renmin Han, Zheng Han, Ting Yu, Guojun Li

## Abstract

Accurate centromere annotation is essential for elucidating chromosomal stability, gene regulation, and the complexities of genome architecture. However, existing methods are often constrained by their dependence on prior knowledge and their limited applicability across diverse genomic contexts. In this study, we present Centromere Annotator (centroAnno), a novel de novo algorithm tailored for the precise annotation of centromeres and tandem repeats directly within complex genomes, assemblies, centromeric sequences, or raw sequencing data. Through extensive evaluations on both simulated and real-world datasets, centroAnno consistently outperforms existing tools in annotation accuracy. Additionally, centroAnno significantly enhances efficiency, achieving annotation speeds 12 to 44 times faster than other methods when applied to human centromeric regions.

## Background

Understanding the structure of the centromere is essential for assembling centromeres [1], studying diseases [2], studying gene expression regulation [3], and comprehending cell division [4]. Accurately annotating the centromere is a fundamental step in analyzing its structure [5]. The centromere region is rich in satellite DNA sequences, which are characterized by long tandem repeats. These sequences often appear as patterns like *m*-*m*-*m*-*m*…, where *m* represents a repeating unit known as a monomer. The length of these monomers varies among different organisms. For example, vthe monomer length in the human centromere region is approximately 171 base pairs (bp), whereas it is 41 bp in chickens [6]. In some organisms, the centromere region may contain multiple types of monomers, with identities ranging from 50% to 90%. These monomers repeat in an organized manner to form higher-order tandem repeat units (HORs) [7]. For instance, in the X chromosome of the human genome, the satellite DNA sequence follows a pattern like *m*_1_-*m*_2_-…-*m*_11_-*m*_12_-*m*_1_-*m*_2_…-*m*_11_-*m*_12_…, where *m*_1_-*m*_2_-…-*m*_11_-*m*_12_ repeats multiple times, forming a HOR with 12 monomers. A key goal of centromere annotation is to identify monomers and HORs within the centromeric region.

A key challenge in centromere annotation is repeat annotation, and a series of tools have been designed for this classic problem, e.g., Tandem Repeat Finder (TRF) [8], RepeatExplorer2 [9], and TAREAN [10]. TRF can *de novo* extract repeats from tandem repeat sequences while RepeatExplorer2 uses a clustering approach for repeat annotation. TAREAN builds graphs based on raw sequencing data to identify circle structures as repeats. Although these tools perform well for repeat annotation, none of them support the analysis of HORs.

A range of tools specifically addresses the analysis of HOR in centromeric regions, such as Alpha-CENTAURI [11], CentromereArchitect [5], HORmon [12], and HiCAT [13]. While these tools can automate the analysis of HORs and replace manual work, they require known monomer templates or location information of centromere in genome, thus limiting their applicability to centromere structure analysis in some non-model organisms. Additionally, tools such as CentromereArchitect, HORmon, and HiCAT operate on accurate centromeric assemblies, further limiting their usefulness. To address these challenges, TRASH [14] and Satellite Repeat Finder (SRF) [15] were proposed. TRASH can *de novo* analyze monomers from centromeric sequences and infer HORs. However, TRASH requires the user to manually select monomers from an inferred monomer set as templates for HOR analysis. More than just affect automation, this manual selection may not always produce accurate HOR results. SRF reconstructs satellite repeats/HORs from accurate sequencing data, such as pacbio High-Fidelity (HiFi) data and next-generation sequencing data [16]. To obtain annotation results, it is necessary to map the reconstructed tendem repeats/HORs back to the source sequence. However, aligning tandem repeat sequences is inherently a challenging task, which could result in incomplete annotation outcomes.

To overcome these limitations, we propose a novel automated centromere annotation tool named centroAnno. CentroAnno can *de novo* derive monomers/tendem repeats and HORs from a genome/assembly/centromeric sequence/noisy sequencing read without requiring prior information such as monomer templates or knowledge of whether the sequence is a tandem repeat or centromeric sequence. Additionally, centroAnno can directly recruit potential tandem repeat sequences from noisy sequencing data and analyze their structure. Extensive experiments have demonstrated that centroAnno outperforms existing tools in accurately analyzing centromere regions across multiple species, with analysis speeds up to 12 to 44 times faster than other tools. These advancements provide crucial support for constructing centromere landscapes across multiple individuals within a species.

## Results

### Overview of centroAnno

CentroAnno is designed to process FASTA (or FASTQ) files containing multiple sequences as input and does not require a priori information such as monomers (refer to Figure 1a). Initially, centroAnno employs a heuristic method to assess if a sequence contains tandem repeats. This involves conducting k-mer counting on the sequence, with an option for users to utilize homopolymer compression technology (HPC) to enhance efficiency and filter out potential sequencing errors (refer to Figure 1b). Following k-mer counting, centroAnno calculates the ratio of repeated k-mers. If this ratio surpasses a specified threshold, the sequence is identified as a tandem repeat sequence or contain tandem repeat regions, prompting further analysis (refer to “Methods”).

**Figure 1.**
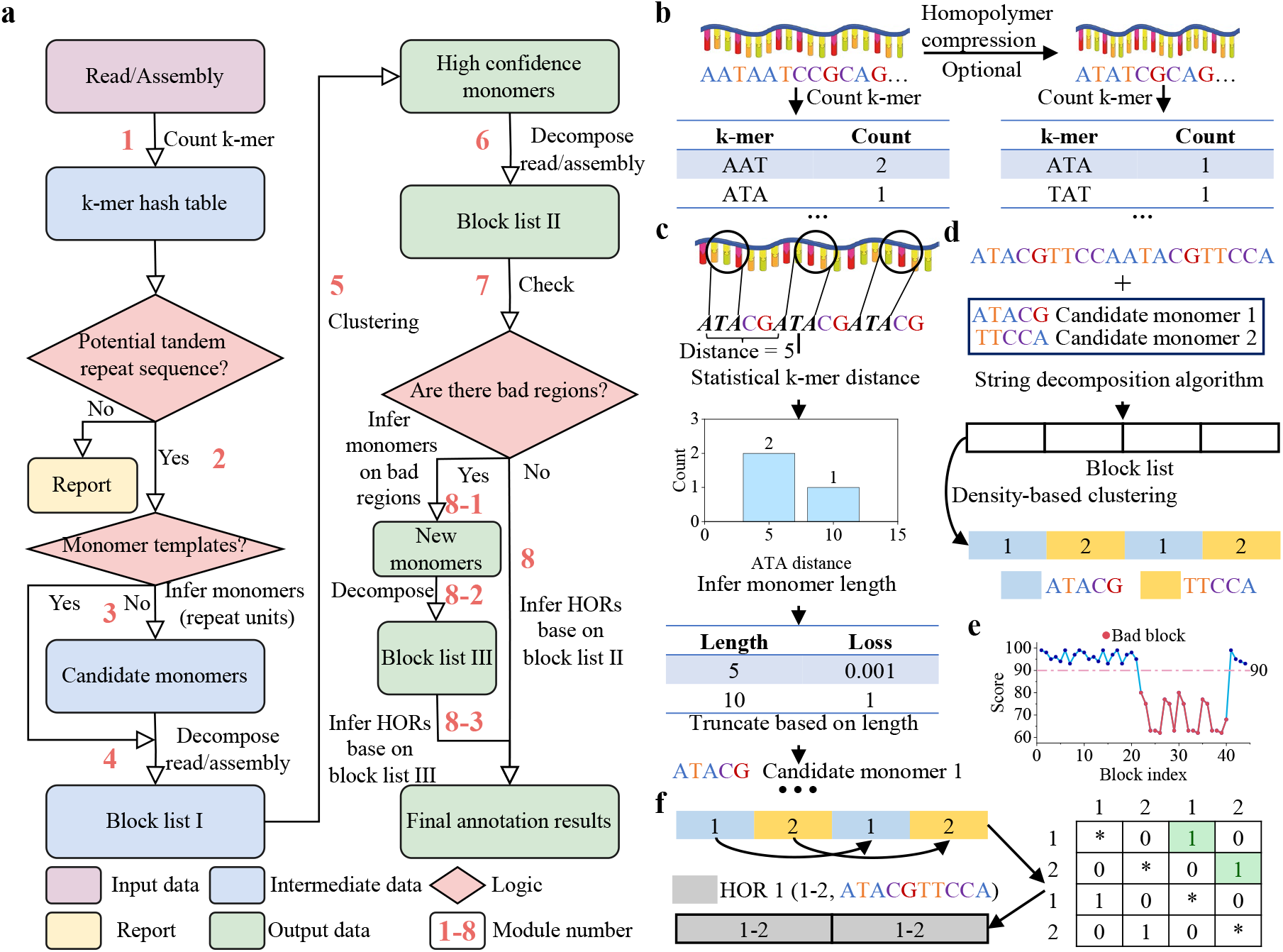
The general workflow of centroAnno. **a**. CentroAnno’s workflow diagram. **b**. Two methods for counting k-mers. **c**. The process of inferring candidate monomers. The k-mer distance refers to the number of bases between two identical k-mers on a linear sequence. **d**. The process of inferring high-confidence monomers from candidate monomers involves using the same string decomposition algorithm as StringDecomposer [16]. Using edit distance [17] as a metric, a density-based clustering algorithm is used to cluster all blocks, and a multiple sequence alignment algorithm is used to obtain the consensus sequence of each cluster as a high-confidence monomer. **e**. Bad blocks in the Block list. These bad blocks form bad regions. **f**. The process of inferring HORs. First, an adjacency matrix about the block list is constructed. “1” implies a “hit”, and HORs are mainly determined from the diagonal containing the most “hits”.

Subsequently, centroAnno selects candidate monomers (repeat units) based on k-mer distance (refer to Figure 1c). These candidate monomers undergo decomposition using a string decomposition algorithm [16] to generate a block list I (refer to “Methods”). Then, centroAnno applies a density-based clustering method [18] to cluster these blocks, resulting in the identification of consensus sequences for each cluster. These consensus sequences are considered high-confidence monomers. CentroAnno utilizes these high-confidence monomers for another round of sequence decomposition to obtain block list II (refer to Figure 1d, “Methods”).

Additionally, centroAnno assesses the quality of each block based on its score (refer to Figure 1e). Here, the score denotes the identity between the monomer template and the block, also known as RepBlock-MonoTemp identity (refer to “Methods”). Poorly scored regions indicate incomplete monomer sets, prompting centroAnno to infer potential monomers in these regions and perform another round of sequence decomposition to obtain block list III (refer to “Methods”). Finally, centroAnno infers HORs based on the final block list (refer to Figure 1f, “Methods”). On the other hand, we developed an automated pipeline with centroAnno as the core to directly analyze tandem repeat units and HORs from raw sequencing data (refer to “Methods”).

### Benchmarking of centroAnno

We assessed the performance of centroAnno using both simulated and real datasets. The length of the monomer within the centromeric region of the genome typically ranged from 100 to 200 bp [14]. For instance, in humans, the monomer length was approximately 171 bp [19], while in rice, there were two monomers with lengths of approximately 155 bp and 165 bp [20]. Some species had even shorter monomers, such as chickens, with a length of 41 bp [6]. To simulate data, we utilized both human and chicken monomer templates. Initially, we generated 10 diversified monomer templates by introducing random mutations of 10%, 15% of the bases in each raw monomer template. These diversified templates were concatenated to form a HOR. Subsequently, the HOR was duplicated 500 times to mimic a typical centromeric region. Finally, centromeric sequences were generated by randomly mutating 0.5%, 1%, or 1.5% of the bases within each centromeric region. This process was repeated three times, resulting in a total of 36 (2 × 3 × 3 × 2) simulated datasets. To evaluate the performance of centroAnno in annotating tandem repeat units, we also simulated a genomic sequence with a length of 100,213,216 bp. This sequence includes 50 tandem repeat regions, each ranging from 50,000 bp to 1,000,000 bp in length and containing tandem repeat units of 40 bp to 200 bp. All scripts for generating the simulated sequences are available at https://github.com/junhaiqi/centroAnno/tree/main/script/simulation. Our real datasets were primarily sourced from the centromere regions of various species, and including raw sequencing data of humans from Oxford Nanopore Technologies (ONT) and pacbio HiFi (see “Data availability”).

We compared centroAnno with state-of-the-art methods such as HORmon, HiCAT, TRF, SRF, and TRASH. HORmon and HiCAT are specialized tools for analyzing centromere structures, both of which require known centromere sequences and monomer templates as inputs. To ensure a fair comparison with these two methods, centroAnno also uses centromere sequences as input, but a monomer template is not mandatory. One of the unique features of centroAnno is its ability to directly analyze HORs within the genome without the prerequisite of centromere sequences. Therefore, we also evaluated the performance of centroAnno in directly analyzing HORs within the genome. Additionally, SRF, TRF, TRASH, and centroAnno all support the de novo inference and annotation of tandem repeat units within sequences, and we compared centroAnno with these tools in this regard. Given our knowledge of the ground truth in the simulated dataset, evaluation metrics included the consistency between the inferred monomer length and the ground truth, as well as the consistency between the inferred HORs and the ground truth. When evaluating all methods on real datasets, our assessment considered the consistency of the repeat (monomer) length with reported information and the consistency of the most frequent HORs inferred by all methods with the reported information. The experimental environment and command lines are provided in Supplementary Note 1.

### CentroAnno outperforms other centromere analysis tools on simulated datasets

We first compared centroAnno, TRF, SRF, and TRASH on simulated genomic sequences. Except for SRF, the other tools successfully annotated the 50 tandem repeat regions (Figure 2a, Supplemental Table S1). This may be because SRF’s annotation process requires mapping the reconstructed tandem repeat monomers back to the source sequence, which can lead to missing some potential tandem repeat regions. Both centroAnno and SRF accurately inferred tandem repeat units, but some of the tandem repeat units inferred by TRASH were consistently off by ∼ 1 bp compared to the ground truth (Supplemental Table S1). TRF, on the other hand, misclassified subsequences of tandem repeat units as complete tandem repeat units (Supplemental Table S1). In terms of speed, centroAnno, TRF, and SRF were all able to complete the analysis within 40 minutes, while TRASH took approximately 3 hours.

**Figure 2.**
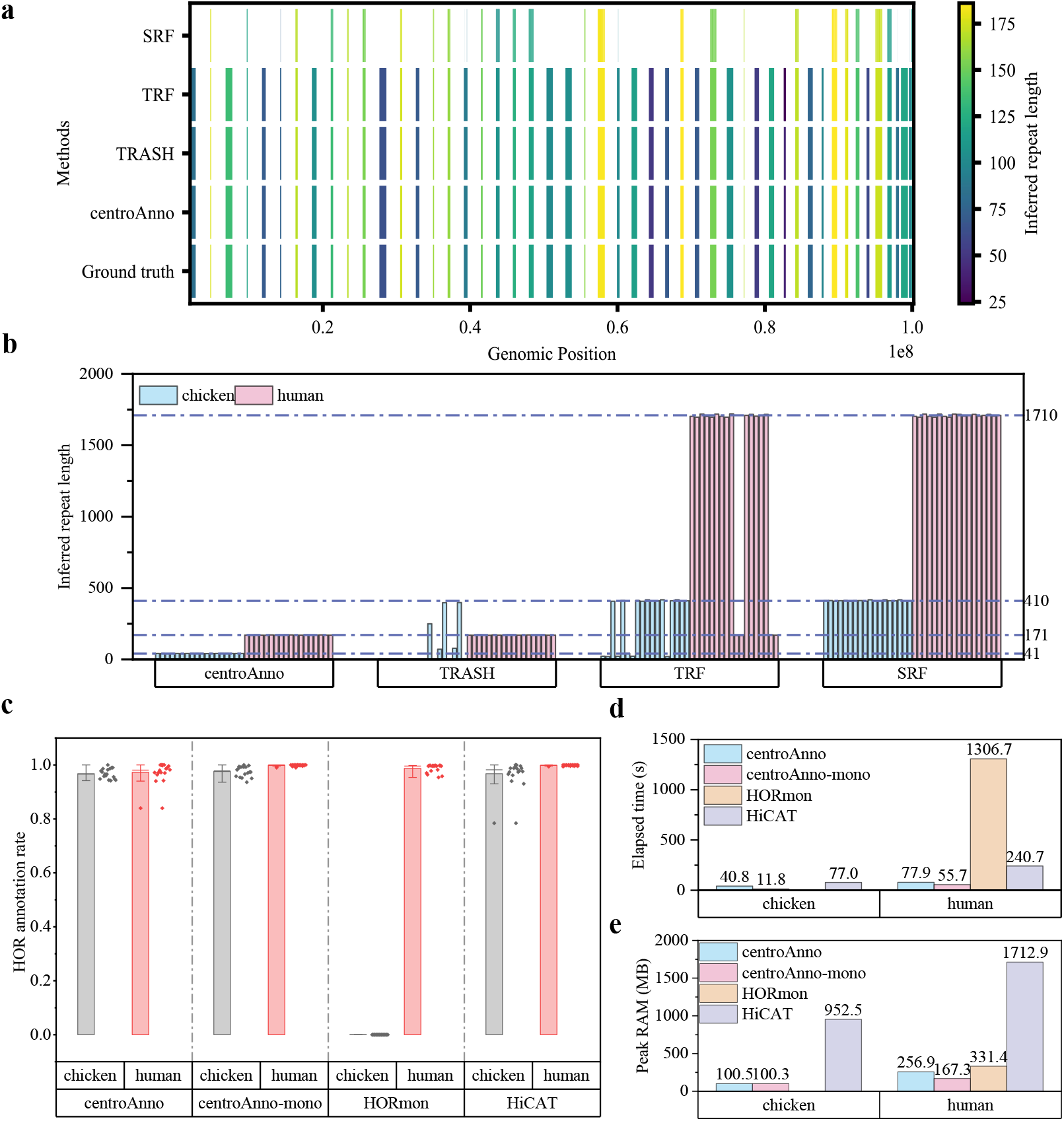
Comparison of centromere analysis methods on the simulated datasets. **a**. Distribution of annotated regions on the simulated genome for all methods. **b**. Repeat unit lengths inferred by all methods on 36 simulated centromere sequences. **c**. Length distribution of HORs inferred by centroAnno, HORmon, and HiCAT on 36 simulated centromere sequences. **d**. Runtime of centromere analysis methods for HOR annotation across all simulated datasets. The figure shows the average running time of each method on all datasets. **e**. Memory usage of centromere analysis methods for HOR annotation across all simulated datasets.

We further compared centroAnno, TRF, SRF, and TRASH on 36 simulated sequences containing HORs (Figure 2b). The simulated sequences of chicken (human) had tandem repeat unit lengths of approximately 41 bp (171 bp), with HOR lengths of about 410 bp (1710 bp). centroAnno inferred the correct repeat lengths across all sequences, whereas TRASH failed to detect tandem repeat units in some simulated chicken HOR sequences (Supplemental Table S2) and in certain datasets incorrectly inferred tandem repeat units as HORs. TRF also inferred HORs in most datasets rather than the correct tandem repeat units, detecting the correct tandem repeat units in only a few datasets. SRF reconstructed HORs in these HOR sequences, consistent with their original report that SRF can detect HORs in HOR regions but cannot further resolve the tandem repeat units comprising the HOR.

Additionally, we compared centroAnno, HORmon, and HiCAT on these simulated sequences containing HORs, and all three tools were able to infer the monomer composition of the HORs. When template information (referred to as centroAnno-mono) is provided, it outperforms other tools on most datasets. Without template information, centroAnno performs comparably to other template-dependent methods. The HOR annotation rate was calculated as 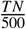, where *TN* represented the number of annotated HORs with a length of 10, and 500 corresponded to the correct number of HORs. The length of the HOR referred to the number of monomers contained. For example, a HOR of length 10 was represented by a 10-monomer HOR. In Figure 2c, HORmon failed to annotate simulated datasets of chicken, and HiCAT also had an annotation rate lower than 0.8 for some chicken datasets. This may have been due to the shorter monomer length of chickens, making HOR identification more challenging. On the other hand, centroAnno demonstrated efficiency and memory advantages. Compared with HORmon and HiCAT, centroAnno completed the analysis 0.9 to 15.9 times faster (Figure 2d) and used 0.2 to 5.6 times less memory (Figure 2e). In addition, centroAnno-mono completed the analysis 5.5 to 23 times faster (Figure 2h) and used 0.9 to 8.5 times less memory.

### CentroAnno outperforms other centromere analysis tools on human CHM13-T2T genome (v2.0)

We obtained the human CHM13-T2T genome (v2.0) [21] and isolated centromeric regions from all chromosomes using SAMtools [22] (see Supplementary Note 2 for details). The performance of centroAnno was compared with other centromere analysis tools on these regions. One of the unique features of centroAnno is its ability to directly annotate centromere regions and centromere-like regions containing tandem repeats within genomic sequences, without requiring prior knowledge of the centromere’s location in the genome or whether the sequence belongs to a centromere. On these genomic sequences, centroAnno not only inferred the 171 bp monomer but also identified common tandem repeat units. For instance, in the telomeric regions, centroAnno inferred (AACCCT)n repeats. In the centromeric regions, it inferred ∼ 42 bp Human Satellite 1 (HSat1) and ∼ 68 bp βsatellite (βSat) [19]. The inferred HORs were consistent with the annotations on centromere regions, these annotation results can be accessed at [23].

We denoted the centromeric region of the Mth chromosome as CENM. For example, CEN1 represented the centromeric region of chromosome 1. Initially, we assessed centroAnno’s repeat annotation performance. Since neither HiCAT nor HORmon supported repeat inference and the results of simulation experiments show that TRF and SRF recognize HOR as a tandem repeat unit, we only compared the repeat annotation performance of centroAnno and TRASH. The repeat annotation rates of centroAnno across all centromeric regions exceeded 0.9 (Figure 3a), while TRASH’s annotation rate on CEN15 was below 0.8. Moreover, centroAnno accurately inferred repeat lengths, mostly aligning with the ground truth of 171bp (Figure 3b). In contrast, TRASH misinterpreted repeat lengths on CEN2, CEN9, CEN15, and CEN21, thereby affecting the annotation rate (Figure 3a, b).

**Figure 3.**
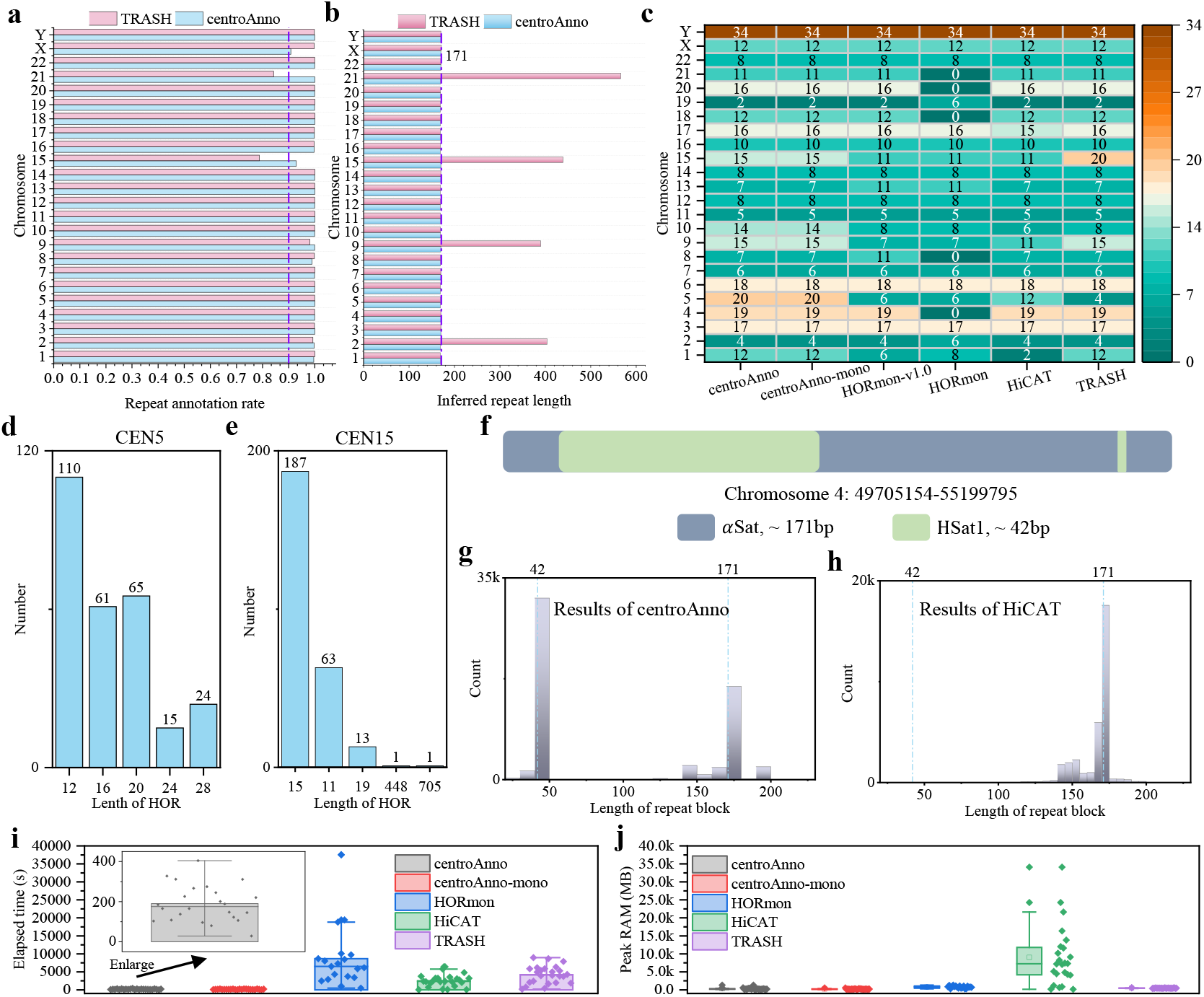
Comparison of centromere analysis methods on human CHM13-T2T genome (v2.0). **a**. Repeat annotation rates of centroAnno and TRASH. **b**. Inferred repeat length of centroAnno and TRASH. **c**. Most frequent lengths of HOR inferred by centromere analysis methods. “0” implies that the analysis failed. HORmon-v1.0 denotes the HOR annotation result of HORmon in the human CHM13-T2T genome (v1.0), obtained from [12]. TRASH’s HOR annotation results are obtained from [14]. **d**. Actual length distribution of the most frequent HORs inferred by HiCAT for chromosome 5. For convenience, only the top five HORs were utilized in the drawing process. **e**. Actual length distribution of the most frequent HORs inferred by HiCAT for chromosome 15. For convenience, only the top five HORs were utilized in the drawing process. **f**. A centromeric region contains αSatellite (αSat) and HSat1. **g**. The repeat (monomer) length distribution inferred by centroAnno on a centromeric region containing αSat and HSat1. **h**. The repeat length distribution inferred by HiCAT on a centromeric region containing αSat and HSat1. **i**. Runtime distribution of methods for centromere analysis of human CHM13-T2T genome. **j**. Memory usage distribution of methods for centromere analysis of human CHM13-T2T genome.

Further analysis focused on centroAnno’s performance in annotating HORs (Figure 3c). While all tools generally provided consistent results across most chromosomes, discrepancies arose in HOR length inference for certain chromosomes. The HOR length inferred by centroAnno differed somewhat on CEN1, CEN5, CEN9, CEN10, and CEN15 compared to other tools. For CEN1, HiCAT inferred a predominant HOR length of 2, while we found that the 12-monomer HOR identified by centroAnno is actually a tandem array of 2-monomer HORs. Similarly, most 8-monomer HORs inferred by HORmon, such as the “A-B-A-B-A-B-A-B” pattern, are also tandem arrays of 2-monomer HORs. centroAnno’s selection of the 12-monomer HOR as the most frequent HOR likely ensures more consistent and continuous annotation. Notably, the most frequent HOR lengths inferred by HiCAT may not always represent the actual HOR length. For example, HiCAT considered the HOR labeled 1–2-3–4-5–6-1–7-(1–2-3–8), which had the same length as 1–2-3–4-5–6-1–7-(1–2-3–8)×3, both were 12, where the numbers represented monomer labels. This was because 1–2-3–4-5–6-1–7-(1–2-3–8)×3 was considered a variation of 1–2-3–4-5–6-1–7-(1–2-3–8). The most common HOR lengths inferred by HiCAT on CEN5 and CEN15 were 12 and 11, respectively, and we determined the actual length distribution of these HORs (Figure 3d, e). For CEN15, HiCAT inferred the actual length of the most frequent HOR to be 15, which was consistent with centroAnno’s inference. On the other hand, centroAnno inferred the most frequent HOR length to be 20 on CEN5, which ranked third among HiCAT’s inferences. On CEN10, TRASH and HORmon provided consistent inferences, while centroAnno and HiCAT yielded inconsistent results. In the results of HORmon, an 8-monomer HOR was followed by a 6-monomer HOR. The HOR inferred by centroAnno may have been the concatenation of these two HORs to ensure more continuous results (Supplementary Note 3). The differences between centroAnno and other tools on CEN9 is due to a similar reason: the 15-monomer HOR identified by centroAnno is actually a tandem array of an 11-monomer HOR and a 4-monomer HOR [15].

Sometimes, we encountered situations where we could not obtain a pure HOR region, and it contained unexpected repeats, such as HSat1 (Figure 3f). In such cases, accurately identifying these repeats became crucial. Notably, centroAnno successfully detected all repeat types in this centromeric region, whereas HORmon was unable to process it effectively, and HiCAT missed the HSat1 repeat (Figure 3g).

CentroAnno completed centromeric region analysis in just 190s on average (Figure 3i, Supplemental Table S3), making it 12 to 44 times faster than other methods. Additionally, centroAnno required an average of only 0.3GB of memory (Figure 3j, Supplemental Table S4), which was 0.03 to 0.6 times the memory usage of other methods. Moreover, when template information was provided, centroAnno-mono outperformed centroAnno in terms of both time and memory efficiency, being on average 30s faster and consuming an average of 0.07GB less memory.

### CentroAnno efficiently analyzes centromere structures across multiple species

In addition to the human genome, we also collected the T2T genomes of four species, namely *Arabidopsis thaliana* [24], rice [20], grape [25], and the chromosome Y of fish [26]. Based on the reported centromere positions, we extracted the centromeric regions of these genomes for analysis (Supplementary Note 2). The length of the monomer in the centromeric region of these species ranged from 100bp to 200bp. In particular, the length of the fish monomer was 524bp (Figure 4a). Current centromere analysis tools typically decomposed sequences into block arrays. Each block was a subsequence of the centromere DNA sequence and exhibited high identity to the monomer template. We recorded this identity as RepBlock-MonoTemp identity. The RepBlock-MonoTemp identity of centroAnno was defined as 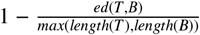. Here, *ed*(*T, B*) represented the edit distance between template *T* and block *B*, and *length*(*) represented the sequence length.

**Figure 4.**
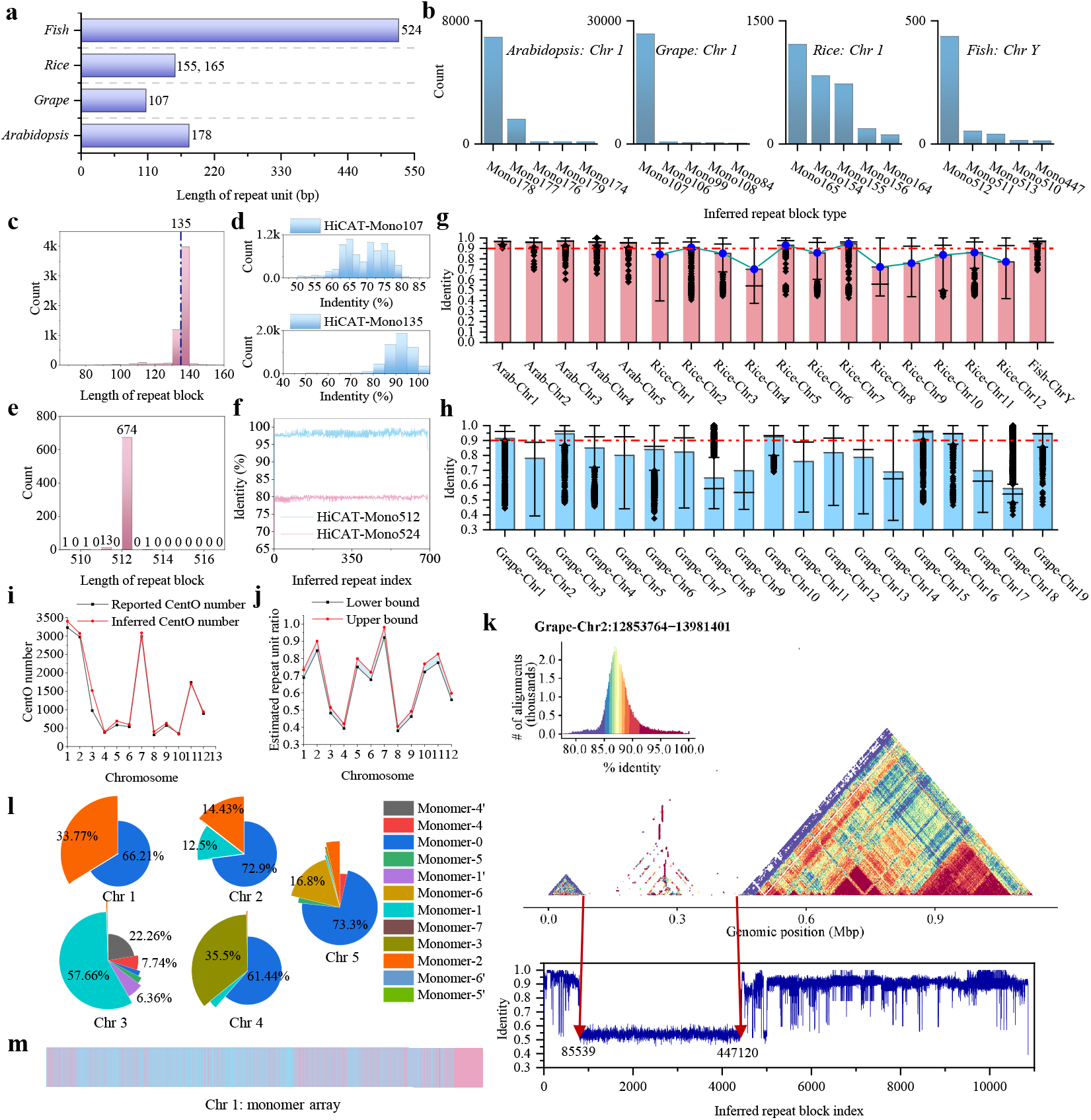
Centromere structure analysis in 4 species. **a**. Reported monomer lengths for 4 species. **b**. Monomer length distribution by centroAnno. “MonoM” represents the monomer of length M inferred by centroAnno. **c**. Length distribution of repeats on Grape’s chromosome 3 by centroAnno. **d**. RepBlock-MonoTemp identity distribution on Grape’s chromosome 3 by HiCAT using reported (Mono107) and inferred (Mono135) monomer templates by centroAnno. **e**. Length distribution of repeats on fish’s chromosome Y by TRASH. **f**. RepBlock-MonoTemp identity distribution on fish’s chromosome Y by HiCAT using reported (Mono512) and inferred (Mono524) monomer templates by centroAnno. **g**. RepBlock-MonoTemp identity distribution of three species by centroAnno. Average values for 12 rice chromosomes shown as a line chart. **h**. RepBlock-MonoTemp identity distribution of grape by centroAnno. **i**. CentO count in rice reported by [20] and inferred by centroAnno. **j**. Estimated repeat unit ratio of rice. **k**. Heatmap of grape chromosome 2 centromeric region (above), and RepBlock-MonoTemp identity distribution by centroAnno (below). **l**. Monomer proportion in *Arabidopsis thaliana* chromosomes. Monomer-x represented as template x, Monomer-x’ as reverse complement. **m**. Monomer array of *Arabidopsis thaliana* chromosome 1. Different colors in the monomer array represent different monomers.

We first used centroAnno to infer the monomers in chromosome 1 and chromosome Y of these species, and the inferred results were consistent with those reported (Figure 4b). After that, we applied centroAnno to all chromosomes. It is worth noting that we obtained monomer lengths on chromosome 3 of grape and chromosome Y of fish that were inconsistent with previous reports. The monomer length inferred by centroAnno on grape chromosome 3 was 135 bp instead of 107 bp (Figure 4c), and TRASH also inferred a 135 bp monomer (Supplemental Figure S1). We inputted these two lengths of monomer as templates into HiCAT. The results showed that the RepBlock-MonoTemp identity generated by the 107bp monomer template was concentrated in 60% - 80%, while the RepBlock-MonoTemp identity generated by the 135bp monomer template was concentrated in 80% - 100%, which implied that the monomer length of grape chromosome 3 was more likely to be 135bp (Figure 4d). In addition, the monomer length inferred by centroAnno on the chromosome Y of fish was 512bp instead of 524bp, which was also supported by TRASH and HiCAT (Figure 4e, f).

We employed centroAnno to analyze the RepBlock-MonoTemp identity distribution of chromosomes across all species (Figure 4g, h). Our analysis revealed that it exhibited notably high RepBlock-MonoTemp identity on *Ara-bidopsis thaliana* and fish, while demonstrating values less than 0.9 on specific chromosomes of rice and grape. Additionally, [20] provided the number of rice monomers (CentOs) in the centromeric region of each rice chromosome (Supplemental Table S5). We identified each high-confidence block (RepBlock-MonoTemp identity > 0.85) inferred by centroAnno as a potential CentO, and our results closely aligned with [20]’s findings (Figure 4i). We further estimated the tandem repeatability of each centromeric region based on the reported number of CentOs (Figure 4j). Specifically, assuming a CentO length of either 155bp or 165bp, we calculated the lower bound of tandem repeatability as 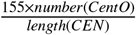 and the upper bound of tandem repeatability as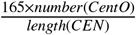, where *number*(*CentO*) represents the count of CentOs and *length*(*CEN*) denotes the length of the centromeric region (Figure 4j, Supplemental Table S5). It is noteworthy that the line charts in Figure 4g and Figure 4j exhibited similar trends, indicating that centroAnno’s RepBlock-MonoTemp identity could effectively estimate tandem repeatability.

The repeat annotation rates of centroAnno and TRASH on all chromosomes, as well as the inferred monomer lengths, were generally consistent, but the length of repeat blocks produced by centroAnno was closer to the length of the monomer template (Supplemental Table S6, Supplemental Figure S1, S2). Additionally, we observed that centromeric regions with lower RepBlock-MonoTemp identity also exhibited lower annotation rates. To delve deeper into these regions, we conducted further analysis using StainedGlass [27], and the resulting heatmap indicated that these centromeric regions displayed lower self-similarity (Supplemental Figure S3). For example, centroAnno identified a region with a very low RepBlock-MonoTemp identity on grape chromosome 2, suggesting that this region may lack self-similarity. The heatmap generated by StainedGlass corroborated this finding (Figure 4k).

We further investigated the distribution of monomers in *Arabidopsis thaliana* and fish due to their high RepBlock-MonoTemp identity. Figure 4l illustrates the distribution of *Arabidopsis thaliana* monomers inferred by centroAnno-mono. We observed that the centromeric region of *Arabidopsis thaliana* typically consisted of 2 to 3 monomers. For instance, an array formed by two monomers ultimately constituted the centromeric region of *Arabidopsis thaliana* chromosome 1 (Figure 4m), which was consistent with previous HiCAT analysis. Additionally, we noted that the centromeric region of the fish chromosome Y mainly comprised one monomer (Supplemental Figure S4), which was also consistent with previous analyses [26].

### CentroAnno is capable of directly analyzing tendem repeats and HORs on noisy sequencing data

When analyzing tandem repeats and HORs from noisy sequencing data, the basic challenge lay in identifying potential reads containing tandem repeats. These k-mers, repeated more than once within a sequence, were referred to as repeated k-mers. An intuitive approach was to look for sequences with a high ratio of repeated k-mers, as such sequences were likely to contain tandem repeats. Experiments also supported this intuition, as shown in Figure 5a and 5b. It could be observed that the majority of k-mers in randomly generated sequences appeared only once, whereas k-mers in centromeric sequences were predominantly repeated k-mers. Despite sequencing errors potentially causing some repeated k-mers to be lost, the overall ratio of repeated k-mers remained high. Consequently, tandem repeat sequences could still be effectively identified based on this ratio of repeated k-mers (Figure 5c).

**Figure 5.**
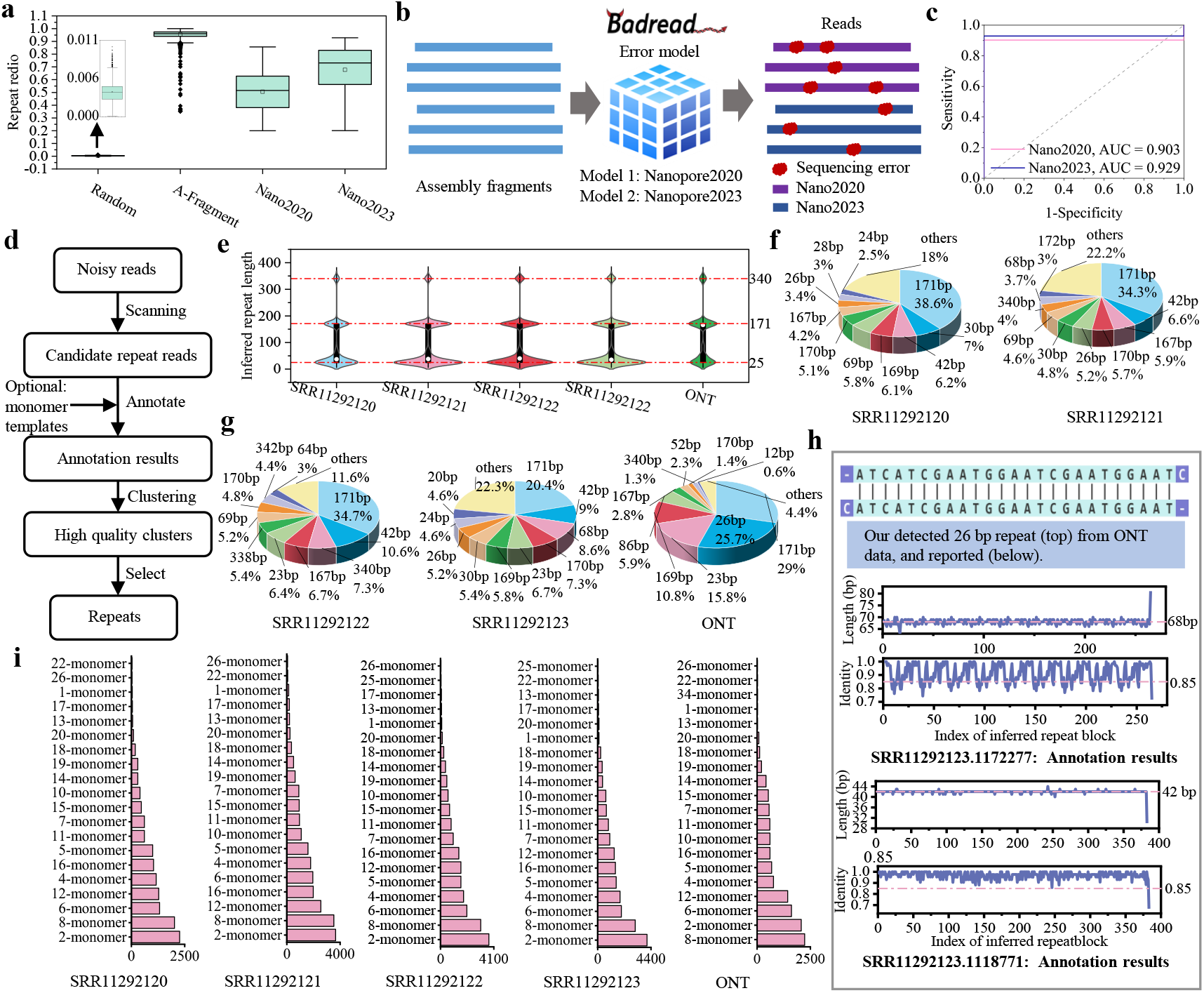
Analysis results of centroAnno on noisy sequencing data. **a**. Ratio of repeated 13-mers for sequences from four datasets. Randomly generate 5,000 sequences with a length of 10,000, and the set of these sequences is called “random”. Randomly truncate 5,000 sequences with a length of 10,000 on human CENX, and the set of these sequences is recorded as “A-Fragment”. The sequencing data generated by two different error models of Badread [28] are denoted as “Nano2020” and “Nano2023” respectively, with “A-Fragment” as the input. **b**. Process for simulating nanopore sequencing data. **c**. Receiver Operating Characteristic (ROC) curve and Area Under the Curve (AUC) value generated by binary classification of sequences from two sequencing datasets, using the ratio of repeated k-mers as the basis. Reads from “Nano2020” and “Nano2023” are used as positive samples, and sequences from “random” are used as negative samples. **d**. A pipeline based on centroAnno for analyzing repeats and HOR directly from noisy sequencing data. **e**. The length distributions of repeats were analyzed by centroAnno across different sequencing datasets. ‘SRR^*^’ represents the accession code of the National Center for Biotechnology Information (NCBI), while ‘ONT’ data can be accessed from https://s3-us-west-2.amazonaws.com/human-pangenomics/T2T/CHM13/nanopore/rel8-guppy-5.0.7/reads.fastq.gz. **f**. The proportions of repeats analyzed by centroAnno on two sequencing data sets. **g**. The proportions of repeats analyzed by centroAnno on three sequencing data sets. **h**. Three typical repeats inferred by centroAnno. “Identity” refers to RepBlock-MonoTemp identity. **i**. HOR distributions inferred by centroAnno on 5 sequencing datasets.

We used centroAnno to analyze repeats directly on HiFi and ONT sequencing data, as detailed in Supplementary Note 4. The entire analysis process was streamlined into a pipeline (refer to Figure 5d and detailed information in “Methods”) to automatically scan and annotate reads containing tandem repeats within the sequencing data. This pipeline also facilitated the discovery of hidden repeats and the HOR structure within the centromeric region. Upon analysis, we observed that the lengths of the majority of the repeats analyzed in these sequencing datasets were clustered around 25 bp and 171 bp, with a smaller number of repeats concentrated around 340 bp in length (refer to Figure 5e). Based on our prior knowledge, we inferred that the 171 bp repeats originated from αSat regions. However, due to sequencing errors, centroAnno incorrectly identified two ∼ 171 bp monomers as a single repeat, leading to the identification of a small number of 340 bp repeats. Furthermore, we conducted an analysis of the composition and proportions of these repeats (see Figure 5f and g, Supplemental Table S7). Our findings revealed that in addition to the ∼ 171 bp repeats, there were also other repeats with relatively higher proportions. These repeats were primarily clustered around lengths of 42 bp, 68 bp, and 26 bp. We conducted a detailed analysis of these three lengths of repeats (refer to Figure 5h). The high-frequency 26 bp repeats we identified have been previously reported by [29]. Moreover, the high-frequency 42 bp repeats and high-frequency 68 bp repeats were found to be repeated in tandem within the reads, corresponding to the reported HSat1 and βSat. Additionally, the pipeline based on centroAnno also analyzed some typical short tandem repeat sequences, such as (TG)n, (CGG)n, (TTCCA)n, etc.

Our pipeline also included support for HOR analysis. Figure 5i illustrated the length distribution of HORs, and we obtained consistent results across different sequencing datasets. Notably, the most frequently observed HORs were the 2-monomer and 8-monomer HORs. This observation could be attributed to the presence of these HOR lengths on multiple chromosomes, as shown in Figure 3c. Furthermore, by combining the analysis results of multiple tools on assemblies (refer to Figure 3c), we observed that the frequently occurring lengths of HORs corresponded to specific chromosomes. For instance, the 6-monomer HOR was prevalent on chromosomes 7 and 10, while the 4-monomer HOR was frequently found on chromosome 2. These findings suggested that the HORs identified by our method in the noisy sequencing dataset aligned well with those inferred from accurate assemblies. This indicated that we could conduct comprehensive centromere structural analysis without assembling the centromere region, significantly reducing the time required for analysis.

## Discussion

CentroAnno is a tool designed for the de novo annotation of tandem repeats and HORs in raw sequencing sequences, genomes, or assemblies, especially centromeric sequences or centromeric regions in the genome. Existing tools either fail to resolve HORs in centromeric regions or necessitate a priori information such as monomer templates or knowledge of whether the sequence is a tandem repeat or centromeric sequence. centroAnno circumvents these limitations by requiring only a user’s input of a single sequence to analyze hidden monomers/tendem repeats and HORs within that sequence. Additionally, centroAnno supports the direct recruitment of potential tandem repeat sequences from raw sequencing data, a capability not currently found in any other tool to our best knowledge.

CentroAnno employs k-mer frequency information to assess whether a sequence exhibits tandem repeat characteristics or includes tendem repeat regions. Furthermore, using a heuristic strategy, we transform the monomer inference problem into an optimization problem, the feasibility of which has been theoretically verified. Through iterative refinement, centroAnno confidently infers monomers, forming the basis for decomposing the tandem repeat sequence into a list of blocks, where each block represents a monomer. centroAnno generates an adjacency matrix based on this block list and derives the HOR from this matrix. Importantly, we developed a centroAnno-centric pipeline for fully automated extraction of repeat sequences and HORs from raw sequencing data.

We evaluated centroAnno using datasets from multiple species. The results demonstrate that centroAnno achieves comparable HOR annotation results to tools using prior information while also inferring monomers more accurately. CentroAnno does not require monomer/tandem repeat unit template information or centromere position information in the genome, this highlights centroAnno’s significance in studying centromeric regions in non-model organisms lacking prior information. Additionally, centroAnno inferred some monomers inconsistent with previous reports, such as those on the fish Y chromosome. Experimental analysis suggests that the monomers inferred by centroAnno may be more reliable, hinting at its potential to discover additional monomers and improve the current monomer library.

Significantly, centroAnno exhibits substantial efficiency and memory advantages, especially when analyzing lengthy centromeric regions. For instance, when analyzing the centromeric region of human chromosome 1, centroAnno proved to be 22 to 35 times faster than other tools (Supplemental Table S3) and used 1.5 to 93 times less memory (Supplemental Table S4). Despite ongoing improvements in algorithm performance and data quality, centromere region assembly remains a significant challenge [30]. Consequently, some centromere studies resort to using raw sequencing data. Our pipeline can directly extract monomers and HORs from such raw data, yielding results consistent with those from accurate assembly. This underscores the vital role of centroAnno in facilitating centromeric landscape studies based on sequencing data from multiple individuals.

## Methods

### The identification of tandem repeat sequences/regions

Tandem repeat sequences consist of one or more repeating units. A natural intuition is that k-mers within tandem repeat sequences occur multiple times. Based on this intuition, we initially assessed whether a sequence was a tandem repeat sequence by analyzing the k-mer distribution. Specifically, for a given sequence, we tallied the occurrences of each k-mer (Figure 5a, Module 1), and k-mers with more than one occurrence were designated as repeated k-mers. The proportion of repeated k-mers among all k-mers was noted as the k-mer repetition rate. When the k-mer repetition rate exceeded a specified threshold, we concluded that the sequence was a candidate tandem repeat sequence. Similarly, we used a sliding window approach to detect potential tandem repeat regions in genome sequences. In contrast, for a non-tandem repeat sequence, its k-mer repetition rate was significantly lower (Figure 5a), making the selection of the threshold relatively straightforward. In our algorithm, the threshold defaults to 0.2. This determination method that utilizes a k-mer counting strategy enabled centroAnno to rapidly screen candidate tandem repeat sequences. For example, centroAnno identified 137,731 candidate tandem repeat sequences from SRR11292123 (containing 1,566,040 reads) in approximately 15 minutes (Supplementary Note 4). CentroAnno then decomposed the sequence into a block list I (Figure 1a, Module 4) and provided a score (RepBlock-MonoTemp identity) for each block. When the mean score of all blocks was greater than a given threshold (default is 85), it was finally judged as a tandem repeat sequence. Additionally, we offered homopolymer compression technology (Figure 1b), which expedited analysis while aiding in monomer (tandem repeat unit) inference, particularly with accurate assembly data (Supplementary Note 5).

### The inference of candidate monomers (tandem repeat units)

When monomer templates are not provided, we employ a heuristic strategy (Figure 1c) to identify candidate monomers: initially, we determine their length based on the distance between two identical k-mers, known as the k-mer distance, and subsequently truncate the sequence between two k-mers. The resulting sequence represents a candidate monomer. This strategy raises two inquiries: 1) What is the likelihood that adjacent monomers share a k-mer? 2) How can we accurately ascertain the true monomer length when faced with multiple k-mer distances? The answer supporting our strategy for the first question is the high probability of two monomers sharing a k-mer. Once we have the expected answer to the first question, we then address the second question, which revolves around avoiding the misidentification of multiple consecutive monomers as a single monomer. Next, we demonstrate that when the sequence is long enough, the probability of two adjacent monomers sharing a k-mer is sufficiently high. Simultaneously, we transform the problem of determining the monomer length into an optimization problem.

From an evolutionary perspective, any monomer evolves from an ancestral monomer, which we denote as *m*. Assuming that the mutation (evolution) rate of each base in *m* is *E*, mutations can occur in three ways: insertion, deletion, and substitution. We refer to the k-mers of *m* as ancestral k-mers, and any k’-mer of a monomer is induced by an ancestral k-mer. For example, if *m* is ATC GAC TAC and a monomer is AT GTC TACG, due to deletion, substitution, and insertion, some ancestral 3-mers evolve into 2-mers, 3-mers, and 4-mers in the monomer: ATC → AT, GAC → GTC, TAC → TACG. If k’-mer and k”-mer, each from a different monomer, evolve from the same ancestral k-mer, we say that k’-mer and k”-mer form a k-pair. We denote a pair of adjacent monomers in a tandem repeat sequence as *m*_1_, *m*_2_, and consider a k-pair regarding *m*_1_, *m*_2_, with their ancestral k-mer denoted as *k*-*mer*_ancestor_. The probability of the “k-mers” (denoted as 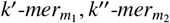) in the k-pair being the same is represented as *P*_*k*_. If 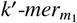 and 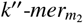 have not evolved (denoted as event *A*), meaning 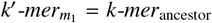 and 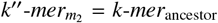, then 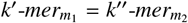 must be true. We denote *P*(*) as the probability of event * occurring, and then *P*(*A*) ≤ *P*_*k*_ holds.

Since the probability of a base mutation in *k*-*mer*_ancestor_ is *E*, and bases are independent of each other, the probability of 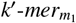 (or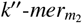) has not evolved is (1 − *E*)^*k*^. Therefore, *P*(*A*) = (1 − *E*)^*k*^ × (1 − *E*)^*k*^ = (1 − *E*)^2*k*^ ≤ *P*_*k*_, and then 1 − *P*(*A*) = 1 − (1 − *E*)^2*k*^ ≥ 1 − *P*_*k*_ holds. Assuming the length of *m* is *L*, there exist 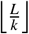 k-mers in *m*, where any two k-mers are non-overlapping and therefore evolve independently. For example, if *m* is ATC GAC TAC, it has 9/3 = 3 non-overlapping 3-mers, namely ATC, GAC, TAC. These 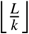 k-mers in *m* evolve to 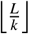 non-overlapping “k-mers” in *m*_1_ and *m*_2_, forming 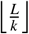 k-pairs that are mutually non-overlapping. The set formed by these k-pairs is denoted as 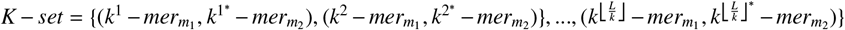. Similarly, in a sequence *S* containing *N* monomers, adjacent monomers form a monomer pair, resulting in 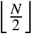 monomer pairs in *S*. Any two monomer pairs are disjoint. The set composed of these monomer pairs is denoted as *KM* − *set*. Consider event *B*: for any k-pair 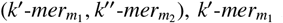 is not equal to 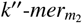. Event *B*^′^: For any k-pair 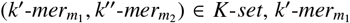 is not equal to 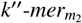. Event *C*: For a sequence containing *N* monomers, considering any monomer pair (*m*_1_, *m*_2_), there is no k-pair 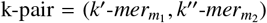, such that 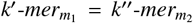. Event *C*^′^: For a sequence containing *N* monomers, considering any monomer pair (*m*_1_, *m*_2_) ∈ *KM* − *set*, there is no k-pair 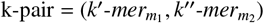, such that 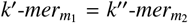. Obviously, event *B*^′^ is a necessary condition for event *B*, and event *C*^′^ is a necessary condition for event *C*, so *P*(*B*^′^) ≥ *P*(*B*) and *P*(*C*^′^) ≥ *P*(*C*). From the above inference, we conclude:

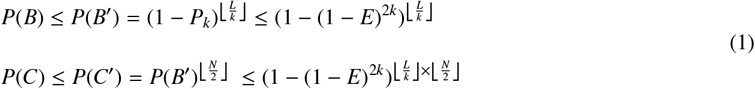

The numerical value of *P*(*C*) is pertinent to our first question. A smaller *P*(*C*) indicates a higher success rate for our heuristic strategy. The above formula provides an upper bound for *P*(*C*). Assuming *N* = 20, mutation rate *E* = 0.15, *L* = 171, and *k* = 10, then *P*(*C*) ≤ 0.0012, suggesting that our heuristic strategy has a high success rate. In addition, if we substitute (1 − *E*)(1 − *e*) for (1 − *E*) in the formula above to account for sequencing error *e*, we can generalize the upper bound of *P*(*C*) in the presence of noisy reads. If *e* = 0.05, then *P*(*C*) ≤ 0.092, further supporting the feasibility of our heuristic strategy.

We have demonstrated that the probability of two adjacent monomers sharing a k-mer is sufficiently high. Next we describe how to determine the true monomer length based on the information of multiple k-mer distances. Given a k-mer, let *PL*_*k*−*mer*_ = [*p*_1_, *p*_2_, …, *p*_*n*_] be the list of positions where it appears in the sequence, with *PL*_*k*−*mer*_ satisfying the condition *p*_1_ < *p*_2_ < … < *p*_*n*_. *PL*_*k*−*mer*_ corresponds to a distance list *D*_*k*−*mer*_ = [*p*_2_ − *p*_1_, *p*_3_ − *p*_2_, …, *p*_*n*_ − *p*_*n*−1_].

Now, consider a positive integer *N* and a deviation *B*. For each *d* ∈ *D*_*k*−*mer*_, if there exists a set {*D*_*i*_ ∈ *D*_*k*−*mer*_ | *D*_*i*_ − *d* < *B, i* = 1, 2, …, *N*}, we define *d* as satisfying the confidence condition *C*_*N*,*B*_, where *N* represents the minimum number of supporting elements for a k-mer distance and *B* indicates the maximum value of supporting elements for that distance. Given a sublist *AD* = [*ad*_1_, *ad*_2_, …, *ad*_*m*_] of the distance list generated by all k-mers, we can model the problem of determining the length of the monomer as the following optimization problem:

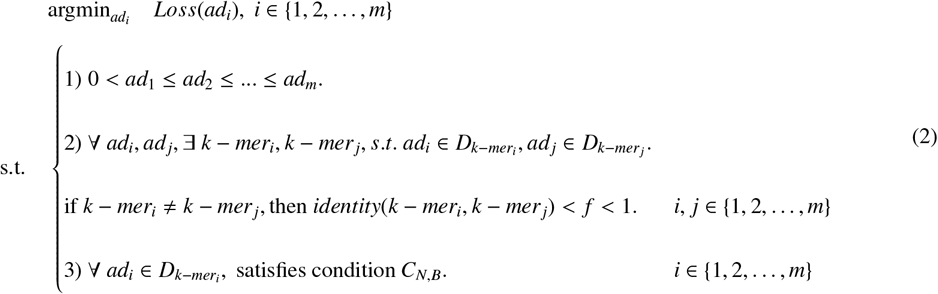

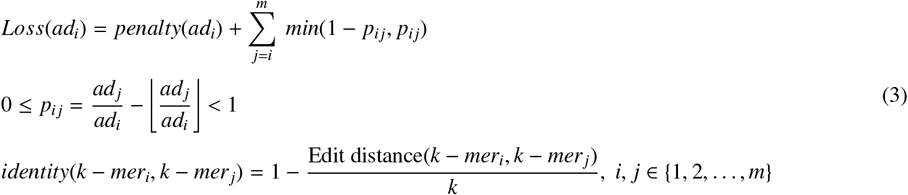

where *Loss*(*ad*_*i*_) measures the loss of *ad*_*i*_ as the repeat length, *identity* measures the similarity between the two sequences, and *f* is a similarity threshold. *penalty*(*ad*_*i*_), equal to *i* − 1, serves as a penalty function, where *penalty*(*ad*_*i*_) represents the number of all elements less than *ad*_*i*_, and min(1 − *p*_*i j*_, *p*_*i j*_) denotes the cost that *ad*_*i*_ is a factor of *ad*_*j*_. We iterate through the distance list starting from the first element, and upon finding a reliable distance, we terminate the iteration and extrapolate this distance as the monomer length. Constraint 1) ensures that a minimum k-mer distance can be identified as the candidate monomer length in this iterative process, while maintaining the utility of *Loss*(*). Similar monomers can be inferred from k-mers with high similarity. Constraint 2) guarantees low redundancy of k-mer set used for inferring the monomer length, achieved through the farthest point sampling algorithm. Constraint 3) ensures an adequate occurrence of k-mer distances in the list, implemented based on *C*_*N*,*B*_. A relatively small and frequent k-mer distance most likely corresponds to the typical length of a monomer, see Supplementary Note 6 for a detailed description of the optimization program.

When the above optimization problem is solved, we can infer the length of the candidate monomer. We collect all k-mers that support this length and the corresponding position information, and obtain candidate monomers through truncation. Furthermore, in order to ensure the efficiency of subsequent sequence decomposition, we use a fast greedy clustering scheme [31] to filter redundant monomers and obtain the final candidate monomer set.

### Inference of high-confidence monomers and decomposition of tandem repeat sequences

We utilize a string decomposition algorithm [16] in conjunction with candidate monomers to break down the tandem repeat sequence into a block list I (Figure 1a, Module 4). Each block in the list is then labeled with its corresponding monomer, indicating that the block has the highest matching score (RepBlock-MonoTemp identity) with the monomer. Subsequently, an edit distance matrix corresponding to the block list is computed. This matrix, coupled with a density-based clustering scheme [18], is employed to cluster the blocks. Following clustering, we calculate the consensus sequence for each cluster using a multiple sequence alignment algorithm, and the consensus sequence serves as a high-confidence monomer. Finally, leveraging all high-confidence monomers and the string decomposition algorithm, we perform another round of sequence decomposition to generate a new block list (block list II in 1a, Module 6). Generally, the scores of blocks in the new list significantly improve compared to the previous block list, indicating enhanced annotation accuracy.

### Further refinement of the decomposition (annotation) results

When not all monomers are inferred, some blocks will receive lower scores (Figure 1e), leading to two scenarios: 1) Multiple consecutive blocks may have low scores, thus forming multiple regions with low scores. 2) Blocks with lower scores may be dispersed throughout the tandem repeat sequence. Upon obtaining the decomposition results, we evaluate the score of each block and identify all blocks with scores below a certain threshold (default threshold is 85). If the total length of these blocks exceeds a specified value (default value is 5000), we conduct further refinement of the decomposition results (Figure 5a, Module 8-1, 8-2, 8-3). In the first scenario, we re-infer monomers for each region, and all newly inferred monomers are included in the high-confidence monomer set. In the second scenario, we cluster these blocks to obtain consensus sequences, which are also added to the high-confidence monomer set. Subsequently, leveraging the new high-confidence monomer set, we employ the string decomposition algorithm to once again decompose the tandem repeat sequence, yielding the final block list (block list III in 1a).

### Inference of HOR

Each element in the block list is assigned a monomer label, and we infer the HOR based on the label information (Figure 1e). Specifically, if a tandem repeat sequence is decomposed into *m*_1_-*m*_2_…*m*_*n*−1_-*m*_*n*_, we regard each *m*_*i*_ as a node, *i* ∈ {0, 1, …, *n* − 1}, and build an adjacency matrix *A*:

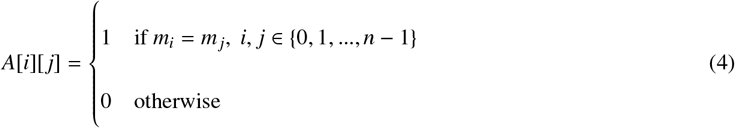

As the adjacency matrix is symmetric, we focus solely on its upper triangular portion. The set of all cells on a diagonal of *A* is termed a wave. *A* has *n* waves, where each *wave*_*i*_ = [*A*[0][*i*], *A*[1][*i* + 1], …, *A*[*n* − *i* − 1][*n* − 1]], for *i* ∈ {0, 1, …, *n* − 1}. The count of elements with a value of 1 in the wave is referred to as the “hit” count. If there exist multiple (greater than 1) consecutive elements with a value of 1 in a wave, we designate these elements as constituting a “hit” region, where the length of the “hit” region corresponds to the number of elements it encompasses.

We identify potential HORs within these “hit” regions. After constructing the adjacency matrix *A*, we first identify the wave (denoted by *wave*_*H*_, *H* ∈ {1, …, *n* − 1}) with the maximum “hit” count, which is considered to contain the most frequently occurring HORs. Then, we search for all “hit” regions in *wave*_*H*_ that meet the following conditions:

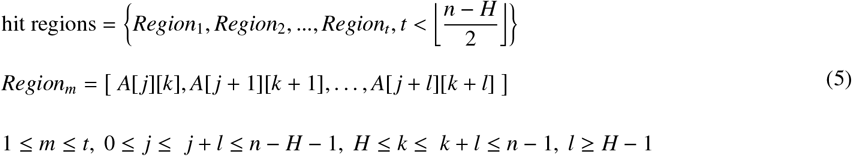

Here, each “hit” region *Region*_*m*_ that meets the conditions determines a HOR region 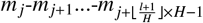, and the typical HOR of this HOR region is *m*_*j*_-*m*_*j*+1_…-*m*_*H*+ *j*−1_ (denoted as *TH*). Subsequently, we create a list *L* = [0, 0, …, 0] of length *n* to record annotation information. When an element in the list is annotated, its value is modified to 1. We iteratively annotate a HOR region and update *L*. Following this, we employ a sliding window strategy combined with the following constraints to identify single HORs not in the HOR region and update *L*:

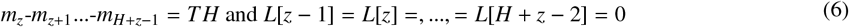

Finally, following the above steps, we iteratively identify and annotate all HOR regions from *wave*_1_, *wave*_2_, …, *wave*_*N*_ (*N* defaults to 50).

### A pipeline for direct analysis of tandem repeats and HORs on noisy sequencing data

We have developed a pipeline centered around centroAnno for direct analysis on noisy sequencing data (Figure 5d). Initially, we identify potential tandem repeat sequences from raw sequencing data. Subsequently, these potential tandem repeat sequences are processed using centroAnno, allowing users to provide monomer (repeat) templates for more precise annotation of tandem repeat sequences containing specific monomers. Following processing, each potential tandem repeat sequence is decomposed into a block list, and we compute the average score of all blocks. If the average score surpasses a specified threshold (default is 85), the corresponding sequence is labeled as a candidate tandem repeat sequence. We determine the most frequently occurring monomer in each candidate tandem repeat sequence to form a monomer set. Finally, the monomer set undergoes clustering using the CD-HIT algorithm [32]. Clusters whose size exceeding a predetermined threshold (default is 5) are designated as high-quality clusters, and their representative sequences are selected as representative repeats. Moreover, in scenarios where monomer templates are provided, we identify sequences that have a monomer similar in length to a given monomer template based on *Loss*(*) in equation (2). These sequences are analyzed using centroAnno-mono to yield more accurate annotation outcomes.

## Data availability

All simulated datasets and results produced by all methods are available in [33], all real assembly data and results produced by all methods are provided in [34], and results of all methods on raw sequencing datasets can be found in [35]. The raw HiFi sequencing data set comes from NCBI with access codes SRR11292120, SRR11292121, SRR11292122, and SRR11292123. The ONT sequencing data set is available at https://github.com/marbl/CHM13.git.

## Code availability

The source code of centroAnno is available at https://github.com/junhaiqi/centroAnno.git, and the pipeline for analysis of raw sequencing data is available at https://github.com/junhaiqi/CentroRepeatAnalyzer.git.

## Acknowledgements

This work was supported by the National Key R&D Program of China with code 2020YFA0712400; the National Natural Science Foundation of China with code 11931008, 12101368, and 61771009, and the Natural Science Foundation of Shandong Province with code ZR2021QA013. The funders had no role in study design, data collection and analysis, decision to publish, or preparation of the manuscript.

## Author contributions

J.Q. conceived and developed the initial version of centroAnno, with input and suggestions from G.L.. J.Q. conducted the benchmarking analyses and contributed to data analysis alongside T.Y. and J.M.. Z.H. provided valuable suggestions for implementing the multiple sequence alignment algorithm. J.Q. drafted the manuscript, which was subsequently revised by T.Y., G.L, and R.H.. All authors reviewed and approved the final version of the manuscript.

G.L. and T.Y. supervised and coordinated the project.

